# Somy evolution in the honey bee infecting trypanosomatid parasite, *Lotmaria passim*

**DOI:** 10.1101/2024.07.12.603340

**Authors:** Lindsey M. Markowitz, Anthony Nearman, Zexuan Zhao, Dawn Boncristiani, Anzhelika Butenko, Luis Miguel de Pablos, Arturo Marin, Guang Xu, Carlos A. Machado, Ryan S. Schwarz, Evan C. Palmer-Young, Jay D. Evans

## Abstract

*Lotmaria passim* is a ubiquitous trypanosomatid parasite of honey bees nestled within the medically important subfamily Leishmaniinae. Although this parasite is associated with honey bee colony losses, the original draft genome—which was completed before its differentiation from the closely related *Crithidia mellificae*—has remained the reference for this species despite lacking improvements from newer methodologies. Here we report the updated sequencing, assembly, and annotation of the BRL type strain (ATCC PRA-422) of *Lotmaria passim*. The nuclear genome assembly has been resolved into 31 complete chromosomes and is paired with an assembled kinetoplast genome consisting of a maxicircle and 30 minicircle sequences. The assembly spans 33.7 Mb and contains very little repetitive content, from which our annotation of both the nuclear assembly and kinetoplast predicted 10,288 protein-coding genes. Analyses of the assembly revealed evidence of a recent chromosomal duplication event within chromosomes 5 and 6 and provides evidence for a high level of aneuploidy in this species, mirroring the genomic flexibility employed by other trypanosomatids as a means of adaptation to different environments. This high-quality reference can therefore provide insights into adaptations of trypanosomatids to the thermally regulated, acidic, and phytochemically rich honey bee hindgut niche, which offers parallels to the challenges faced by other Leishmaniinae during the challenges they undergo within insect vectors, during infection of mammals, and exposure to antiparasitic drugs throughout their multi-host life cycles. This reference will also facilitate investigations of strain-specific genomic polymorphisms, their role in pathogenicity, and the development of treatments for pollinator infection.

## INTRODUCTION

The family Trypanosomatidae (Euglenozoa: Kinetoplastea) is a diverse group of unicellular flagellated parasites that infect plants and many animals, including certain groups of insects (Maslov et al. 2019). Trypanosomatid genomics has focused on insect-vectored, dixenous (two host) taxa in the genera *Trypanosoma* and *Leishmania* due to their clinical relevance as parasites (Albanaz et al. 2023; Kostygov et al. 2024). Consequently, these genera have been the main focus of genomic efforts and have been key to understanding the phylogenetic history, geographic radiation, population dynamics, and therapeutic drug resistance in this family (Lukeš et al. 2018). In contrast, the solely insect-infecting monoxenous trypanosomatids are biodiverse examples of obligate endosymbionts that have received less attention (Lukeš et al. 2018). However, a recent surge in work on these monoxenous relatives has offered several genomic insights into the metabolism, host-specific adaptations for infection, and intraspecific diversity of these species (Kostygov et al. 2024) while also offering clues to the evolution of host shifts and parasitism of humans and other mammals (Flegontov et al. 2016).

Bumblebee (Bombus spp.) and honey bee (Apis spp.) pollinators are particular insects in which monoxenous trypanosomatids seem to thrive (McGhee and Cosgrove 1980; Ryan S. Schwarz et al. 2015) as parasites (see for example: Schmid-Hempel 2001; Ryan S Schwarz et al. 2015; Koch et al. 2017; Gómez-Moracho et al. 2020; MacInnis et al. 2023). Thus, bees are significant and readily accessible models for trypanosomatid-host coevolution and for insight into the way these parasites affect individual fitness and colony success. High quality genomes of the bumblebee trypanosomatids Crithidia bombi and Crithidia expoeki have aided in this effort and provide a basis for studies on population genomics and the identification of genes under selection (Schmid-Hempel et al. 2018). The honey bee parasite Lotmaria passim was first distinguished from its close relative, Crithidia mellificae, in 2015 (Ryan S. Schwarz et al. 2015) and has since been recognized as the dominant trypanosomatid infecting honey bees worldwide (Morimoto et al. 2013; Ravoet et al. 2013; Ravoet et al. 2015; Arismendi et al. 2016; Vejnovic et al. 2018; Xu et al. 2018; Castelli et al. 2019).

The first genome assembly of *Lotmaria passim* was published before its differentiation from *Crithidia mellificae* (Runckel et al. 2014) and has remained the reference for this species despite lacking improvements from newer methodologies and the association of *Lotmaria passim* with honey bee colony loss and reduced individual bee survival in some experiments (Cornman et al. 2012; Gómez-Moracho et al. 2020). Although the absence of a high-quality reference genome has not prevented recent investigations into *Lotmaria passim* physiology and cell biology (Buendía-Abad et al. 2022; Carreira de Paula et al. 2024), transcriptomics (Liu et al. 2020), proteomics and the role specific genes play during infection (Yuan et al. 2024), it does preclude more detailed analysis of important features such as ploidy, repetitive sequences, and an overall gene arrangement.

Here we report the sequencing, chromosome-level assembly, and genome annotation of the BRL type strain of *Lotmaria passim*. Our assembly and annotation match the quality of other recently sequenced insect-specific species (Flegontov et al. 2016; Schmid-Hempel et al. 2018; Albanaz et al. 2023) and improve substantially upon the previously published draft assembly (Runckel et al. 2014). This high-quality reference could provide insights into the adaptations of trypanosomatids to the thermally regulated, acidic, and phytochemically rich honey bee hindgut niche, which offers parallels to the challenges faced by other Leishmaniinae during the challenges they undergo within insect vectors, during infection of mammals, and exposure to antiparasitic drugs throughout their multi-host life cycles. This reference will also facilitate investigations of strain-specific genomic polymorphisms, their role in pathogenicity, and the development of treatments for pollinator infection.

## MATERIALS & METHODS

### Species/Strain Origin

*Lotmaria passim* BRL type strain (ATCC PRA-422) cultures (Schwarz et al. 2015), were grown to early stationary phase in FPFB medium (Salathé et al. 2012) with 10% heat-inactivated fetal bovine serum at 31°C in 25 cm^2^ vented cell culture flasks (Corning Inc., Corning, NY, USA). To generate dense cultures with sufficient DNA for long-read sequencing, ten flasks containing 12 mL of culture were combined before the library preparation steps detailed below. Cultures for sequencing of *Lotmaria passim* BRL (2024) were stored at -80°C until DNA extraction.

Culture stocks were originally isolated from a honey bee gut and expanded in axenic cell culture before cryopreservation in 2013. Derivations obtained via axenic cell culture of this same stock were used for the *Lotmaria passim* BRL (2016) sequencing effort completed by the University of Massachusetts (GenBank Accession ID: GCA_002216525.1). Cultures used for the *Lotmaria passim* BRL (2024) sequencing effort presented herein have since undergone shifts from antibiotic-supplemented Schneider’s medium (Schwarz et al. 2015) to antibiotic-free FPFB medium (Salathé et al. 2012) and have been grown in continuous cell culture for an estimated cumulative timeframe of six months (∼500 generations), in addition to being subjected to several cryopreservation events prior to sequencing.

### Sample Preparation and Sequencing

#### HiFi Library Preparation and Sequencing

High-molecular-weight genomic DNA was extracted for HiFi library preparation from pelleted cultures using the MagAttract^Ⓡ^ HMW DNA Kit (Qiagen, Hilden, Germany). Cell cultures were pelleted at 8000 rpm for 5 minutes and washed with 1 mL of 1X PBS buffer before following all manufacturer instructions. The extracted DNA was pooled and quality-checked using a Qubit 2.0 Fluorometer (Life Technologies, CA, USA) before being stored at -20°C until shipment to the Novogene Corporation (Sacramento, CA, USA) for sequencing.

Long-read DNA sequencing was performed using 5 µg of high-molecular-weight DNA prepared as previously described to generate a PacBio Sequel II HiFi library. Analysis was completed by the Novogene Corporation (Sacramento, CA, USA) on a one-cell PacBio Sequell II (HiFi/CCS mode/cell) machine which generated 20.1 gigabases (Gb) of sequencing data with an average read length of 15,916 and a total length of 20,137,583,129 which were assembled as described below.

#### Hi-C Library Preparation and Sequencing

Genomic DNA was extracted for Hi-C library preparation from freshly pelleted cultures using the Proximo Hi-C Microbe Kit (Phase Genomics, Seattle, WA, USA). Cell cultures were pelleted at 8000 rpm for 5 minutes and washed with 1 mL of 1X PBS buffer before following all manufacturer instructions. The resulting 900 ng product was quality-checked using Qubit 2.0 Fluorometer (Life Technologies, CA, USA), and the resulting Proximo Hi-C Library was stored at - 20°C until shipment to the University of Maryland Institute for Genome Sciences for sequencing.

Short-read DNA sequencing was performed on the Proximo Hi-C Library using Illumina Nextera sequencing (S4 channel 100 bp paired-end reads). This process generated 198.5 Gb of sequencing data (2,250 million read-pairs with an average length of 15,725) assembled as described below.

### Assembly

#### Nuclear Genome Assembly

To assemble the nuclear genome remnant adapter sequences in the HiFi reads were first removed using HiFiAdapterFilt v2.0.0 (Sim et al. 2022) with the default adapter sequence database. Adapter sequences in Hi-C reads were removed using fastp v0.23.4 (Chen 2023). Sequence qualities were checked before and after filtering using fastqc v0.12.1 (Andrews et al. 2012). Haplotype-resolved contigs were assembled from HiFi and Hi-C reads using Hifiasm v0.19.6 (Cheng et al. 2021), and the two assemblies were combined and then scaffolded by Hi-C reads using YaHS v1.1 (Zhou et al. 2023). Manual correction was done in Juicebox v1.11.08 to resolve misassemblies and switch errors (Durand et al. 2016). Finally, chromatin interactions were calculated using HiC-Pro v3.1.0 (Servant et al. 2015) and visualized using HiCPlotter v0.6.6 (Akdemir and Chin 2015).

#### Kinetoplast Genome Assembly

The tandemly repetitive and repeat-free regions of the maxicircle were assembled separately. Reads with partial sequences similar to the cytochrome B gene (accession number: KM980180) were identified using Basic Local Alignment Search Tool (BLAST) v2.14.0 (Altschul et al. 1990). Contigs with BLAST hits were extracted and Tandem Repeats Finder v4.09 was used to locate and trim repeats (Benson 1999). A multiple sequence alignment (MSA) was built from the repeat-free contigs using MAFFT v7.525 (Katoh and Standley 2013) and a consensus sequence was built using the consensusString function from the Biostrings package v2.68.1 in R v4.3.0 (Pagès et al. 2024). All HiFi reads were aligned to the consensus sequence using Minimap2 v2.26 (Li 2018) and the alignment was sorted and indexed using SAMtools v1.6 (Danecek et al. 2021). The consensus sequence was corrected by the alignment using Pilon v1.24 (Walker et al. 2014), and all HiFi reads were aligned to the corrected repeat-free sequences again to check for homoplasy using IGV v2.14.0 (Robinson et al. 2023).

To assemble repetitive regions, HiFi reads that were mapped to both the beginning and the end of the corrected repeat-free sequence were selected, their unmapped regions were extracted, and an MSA was built. The consensus sequence was called and then corrected by the HiFi read alignment and homoplasy was rechecked. The corrected repetitive sequence was concatenated to the end of the repeat-free sequence, and an alignment was built again to check for misassembly and homoplasy.

Minicircle reads were iteratively identified to reconstruct the minicircle sequences. HiFi reads were initially scanned using nhmmscan from HMMER v3.3.2 (Eddy 2011) against a customized profile Hidden Markov Model (HMM) built from the minicircle genome alignment of *Leptomonas pyrrhocoris* (Gerasimov et al. 2021). The top 12 most significant reads were selected and a self-alignment of each read was made to identify the motif of the conserved region (CR): GGGTAGGGGCGTTCACCGA. The two CRs and the downstream sequences were separated using AliView v1.28 (Larsson 2014) and an MSA was built, then a new profile HMM was created from the MSA and the HiFi reads were scanned again. To identify the e-value threshold, the nuclear and kinetoplast genomes were scanned, and the e-value of the top hit (6.30 × 10^-5^) was selected. HiFi reads with an e-value below this threshold were used to build a new profile HMM for the next epoch and the iteration stopped when reads were no longer found. An MSA of the final minicircle read set was built, reads less than 0.1% different were grouped based on the error rates of HiFi sequencing, and consensus sequences were called to avoid duplicates.

### Repeat Analysis

Repeat analysis was performed with RepeatMasker v4.1.6 (Chen 2004) using Euglenozoa as the query species and run with the Dfam database v3.8 (Storer et al. 2021).

### Annotation

Three separate gene-finding pipelines, all using an Augustus-trained model of *Leptomonas pyrrhocoris* (Flegontov et al. 2016), were employed and their non-overlapping results ultimately combined. Transcript evidence for all pipelines was generated by aligning 30 separate publicly available *Lotmaria passim* short read archives (Liu et al. 2020) using Bowtie2 v2.4.2 (very sensitive; end-to-end) (Langmead and Salzberg 2012) to our genome. The resulting SAMtools (Li et al. 2009) converted BAM files were reformatted to GTF files using Cufflinks v2.2.1 (Trapnell et al. 2010), and then further reformatted to either an Augustus hints file or a FASTA file of mRNA transcripts using GffRead v0.12.7 (Pertea and Pertea 2020). The first gene-finding experiment was conducted using Augustus v3.5.0 (Hoff and Stanke 2019) and incorporated the *Leptomonas pyrrhocoris* model and mRNA hints file. The second experiment used the Companion web server (Steinbiss et al. 2016) and incorporated the *Leptomonas pyrrhocoris* model and cufflinks gtf file. The third experiment employed the Maker2 pipeline v3.01.04 (Holt and Yandell 2011) and incorporated the trained model, mRNA transcript evidence, the Euglenozoa RepBase (Bao et al. 2015) protein set for repeat masking, and the complete protein database from TriTrypDB release 67 (Shanmugasundram et al. 2023). Three rounds of Maker2 training were executed: two rounds of Snap model training and one round of Augustus.

The quality of each annotation was assessed using BUSCO 5.7.1 (Seppey et al. 2019) with Euglenozoa OrthoDB v10 proteins (Kriventseva et al. 2019) as a reference database. The first experiment’s annotation produced a complete set of BUSCO proteins with no fragmentation and was then used as the basis for merging the remaining annotation experiments. We subsequently merged the non-overlapping gene predictions from the latter two experiments and assigned gene IDs using AGAT v1.4.0 (Dainat et al. 2020). A final protein set was extracted using GffRead and functional annotations were performed with InterProScan v5.66-98.0 (Jones et al. 2014) using the TIGRFAM (Haft et al. 2013) and PFAM (Mistry et al. 2020) databases with residue annotation disabled, and NCBI BLAST+ (Camacho et al. 2009) using the Uniprot Database (UniProt Consortium 2015) with and e-value cutoff of 1e-5. The resulting functional and GO terms were added to the merged annotation using AGAT. Non-coding RNAs were annotated using cmscan Infernal 1.1.5 (Nawrocki and Eddy 2013) using Rfam-level heuristic filters and skipping all HMM filter stages. Last, we ran pseudogene predictions using Pseudo Finder v1.1.0 (Syberg-Olsen et al. 2022) against the complete protein database on TriTrypDB release 67. We retained only the predicted pseudogenes found in intergenic regions and merged those predictions with the existing annotation as previously described. Final gene annotation statistics were calculated using AGAT.

### Orthology and Phylogenetics

We compared our final protein set to annotations of other Leishmaniinae: *Crithidia acanthocephali*, *Crithidia bombi, Crithidia expoeki*, *Crithidia brevicula*, *Crithidia fasciculata*, *Crithidia mellificae*, *Crithidia thermophilia*, *Leishmania major*, *Lotmaria passim* SF, *Leptomonas pyrrhocoris*, and *Leptomonas seymouri* which originated from (Kostygov et al. 2024), in addition to *Crithidia bombi* SH and *Crithidia expoeki* SH which originated from (Schmid-Hempel et al. 2018), using OrthoFinder v2.5.5 (Emms and Kelly 2019). The results produced a total of 5075 orthogroups with all species present and 2833 of these consisted entirely of single-copy genes. Multisequence alignment, filtering, and trimming were performed using MAFFT v7.525 (Katoh and Standley 2013) and trimAl v1.4.rev15 (Capella-Gutiérrez et al. 2009) as described in Kostygov et al. 2024 (Kostygov et al. 2024). The resulting 986 orthogroups were subset into two separate random selections, without replacement, of 250 orthogroups. Each selection was compared using PhyloBayesMPI v1.9 (Lartillot et al. 2013) under the CATGTR model over two separate chains, until appropriate effect size (1335) and relative difference (0.03) were achieved, at approximately 1700 iterations per chain. The resulting tree file was plotted using the R packages ape v5.7-1 (Paradis and Schliep 2019) and ggtree v3.8.2 (Yu et al. 2017). No differences were found between the trees produced by separate random selections of orthogroups. The initial BUSCO analysis was repeated for all organisms listed here and plotted for comparison.

### Inference of Somy

We compared data for coverage depth and variant distributions per chromosome for several *Lotmaria passim* genome sequencing efforts, including two assemblies for strain BRL: our *Lotmaria passim* BRL (2024) PacBio HiFi reads, *Lotmaria passim* strain BRL Illumina Nextseq reads from 2016 (NCBI Bioproject PRJNA319530; referred to by its ATCC identifier ‘PRA-422’), *Lotmaria passim* SF sequenced by Illumina Nextseq in 2016 (NCBI BioProject PRJNA319529; referred to by its ATCC identifier ‘PRA-403’), and *Lotmaria passim* strains C2 and C3 isolated in 2019 from honey bees in Granada, Spain, as described in (Buendía-Abad et al. 2021) and sequenced by Illumina MiSeq in 2023 (NCBI BioProject PRJNA863431, Ruiz et al., submitted).

Read mapping to our genome assembly was performed using Minimap2 (PacBio HiFi) (Li 2018) and BWA-MEM (all others) (Li 2013) with default settings. Sorting, indexing, and coverage depth of the resulting SAM files was performed with SAMtools v1.13 (Li et al. 2009). Variant calling was then performed using GATK HaplotypeCaller v4.5.0.0 (McKenna et al. 2010). Data cleaning of the resulting VCF files was performed using the R package vcfR v1.15.0 (Knaus and Grünwald 2017) as outlined in Knaus et al. (Knaus and Grünwald 2018).

Chromosome copy number was then inferred based on the three lines of evidence as generated above: median genome coverage, coverage depth by base position within each chromosome, and the distribution of allele frequencies at heterozygous sites. For depth calculations, we mapped raw genomic reads back to the assembly, calculated the median base-level coverage depth for each chromosome, and normalized depth to the median genome coverage. Normalized depths of 0.5x, 1x, 1.5x, and 2x the median were expected for monosomic, disomic, trisomic, and tetrasomic chromosomes, respectively. Patterns of coverage depth across chromosomes were visualized by graphing the base-by-base depth scores. We assessed these plots for a secondary trace at roughly half the depth of the moving average, indicative of heterozygous sites for which half of the reads for the corresponding base were attributed to a disomic chromosome’s alternate haplotype. In contrast, we expected a secondary trace at roughly two-thirds that of the main trace for a trisomic chromosome; multiple secondary traces at one-fourth, one-half, and three-fourths of the main trace for a tetrasomic chromosome; and the absence of any secondary trace to suggest the absence of a distinct haplotype altogether. The distribution of allele frequencies was evaluated by inspection of variant call plots for heterozygous loci. We expected a predominantly 1:1 ratio of divergent alleles in disomic chromosomes (i.e., one allele belonging to each of two haplotypes), a 2:1 ratio for trisomic chromosomes, and a mixture of 1:1 and 3:1 frequencies in tetrasomic chromosomes (Knaus and Grünwald 2018).

### Figures

All visualizations were created using R v4.3.2 Eye Holes (R Development Core Team 2009) and ggplot2 v3.5.0 (Wickham and Wickham 2016) unless otherwise stated.

## RESULTS & DISCUSSION

### Genome Assembly and Annotation

We sequenced, assembled, and annotated the first complete chromosome-level genome of *Lotmaria passim* BRL type strain (ATCC PRA-422; herein referred to as *Lotmaria passim* BRL (2024)) leveraging the combined power of PacBio HiFi sequencing and Hi-C technologies. We achieved separation of the chromosomal haplotigs, where the longer haplotig was included in the principal assembly and the shorter was included in the alternative assembly. The principal nuclear genome assembly comprises 31 chromosomal sequences (Supplementary Figure S1) with a median coverage of 161.0×, while the kinetoplast assembly includes a complete maxicircle and 30 minicircle sequences. The assembly has a total length of 33.7 Mb with a GC content of 54.5% and a contig N_50_ of 1.5 Mb (Table 1, Figure 1). Repetitive elements encompass around 5.8% of the genome (1.9 Mb). Our annotation of the nuclear and kinetoplast genome assemblies identified 10,288 protein-coding genes with a mean gene length of 1,890 bp and 117 tRNA genes with a mean gene length of 75 bp, all of which lack introns. The protein-coding genes span 19.45 Mb (∼57.7%) of the genome, and 439 pseudogenes were predicted, spanning only 320.5 Kb (less than 1% of the genome).

**Figure 1.**
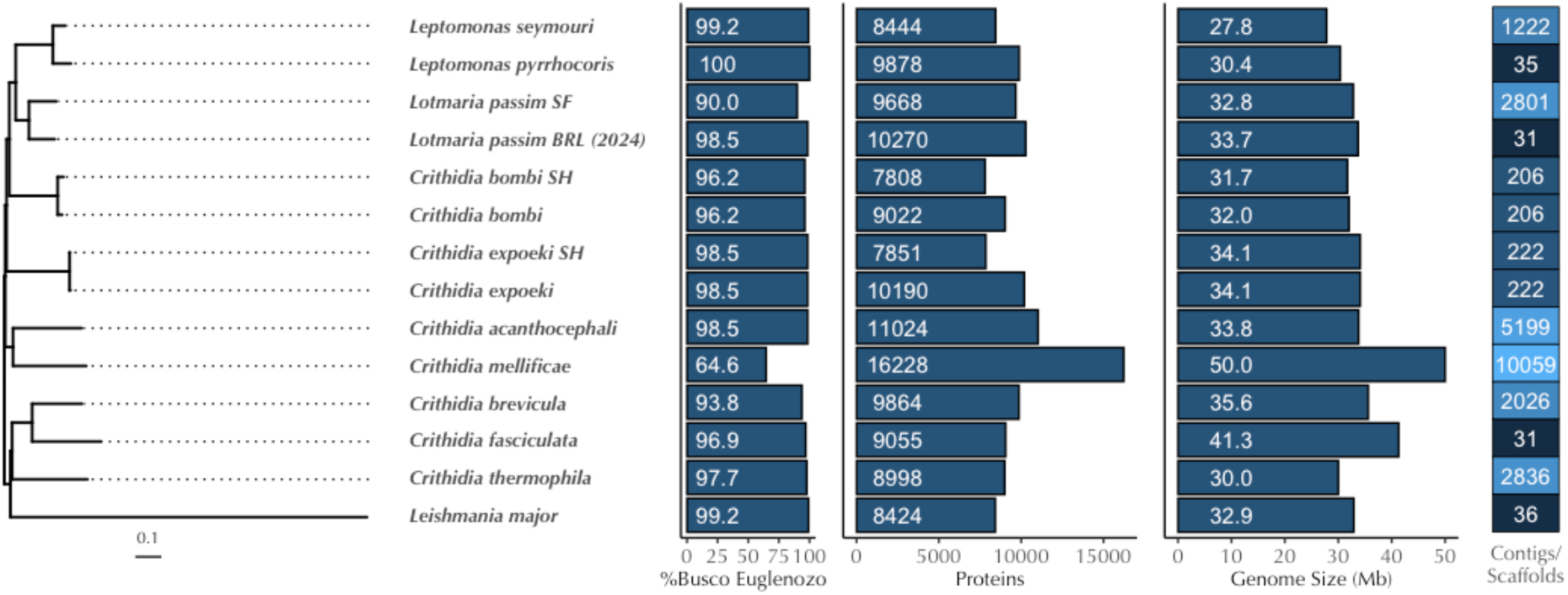
Genome Statistics and Phylogeny of Selected Species Within Leishmaniinae. From left to right: Bayesian phylogenomic tree based on (∼200) single-copy orthologous genes. Columns represent the percentage of complete single-copy BUSCO genes found, the number of annotated proteins, the genome size (Mb), and the number of contigs or scaffolds in species closely related to *Lotmaria passim* BRL. *Lotmaria passim* strain "SF" is based on (Runckel et al. 2014) while *Lotmaria passim* "BRL" refers to the BRL (2024) assembly presented herein. For *Crithidia bombi* and *Crithidia expoeki,* two independent annotations of the same assembly are shown, where the notation "SH" denotes the published annotations in (Schmid-Hempel et al. 2018) and the other branch corresponds to the annotations conducted in (Kostygov et al. 2024)).

**Table 1.** *Lotmaria passim* BRL 422 genome assembly and annotation summary. Mb = Megabases, Kb = kilobases, and bp = base pairs ^†^Half the length of the assembly is contained within contigs of length greater than or equal to the N50. ^‡^The L50 refers to the smallest number of contigs within which half the length of the assembly is contained.

|  |  |
| --- | --- |
| **Table 1.** *Lotmaria passim* BRL 422 genome assembly and annotation summary.<br>Mb = Megabases, Kb = kilobases, and bp = base pairs |  |
| †Half the length of the assembly is contained within contigs of length greater than or equal to the N50. |  |
| ‡The L50 refers to the smallest number of contigs within which half the length of the assembly is contained. |  |
| **Assembly** |  |
| Total genome length, Mb | 33.7 |
| GC content, % | 54.5 |
| Contig N50<sup>†</sup>, Mb | 1.5 |
| Contig L50<sup>‡</sup> | 8 |
| Chromosome number | 31 |
| Repetitive Elements (% of genome) | 1,960,104 bp (5.8%) |
| Maxicircle Assembly Length, Kb | 48.3 |
| Minicircle Assemblies & Lengths | 30 individual assemblies, ranging in length from 360 bp to 1.28 Kb |

| Annotation |  |
| --- | --- |
| Gene number | 10,288 |
| Total gene length, Mb | 19.45 |
| Mean gene length, bp | 1,890 |
| Pseudogene number | 439 |
| Total pseudogene length, Kb | 320.5 |
| Mean pseudogene length, bp | 730 |
| Complete BUSCOs | 130 (100%) |
| Complete, single copy BUSCOs | 128 (98.5%) |
| Complete, duplicated BUSCOs | 2 (1.5%) |

The nuclear genome assembly of *Lotmaria passim* BRL (2024) contains 10,270 proteins (Figure 1), which is within the range of predicted proteins (7,808 to 11,024 (Kostygov et al. 2024)) we have come to expect from other species within the subfamily Leishmaniinae and fit as expected within the Leishmaniinae clade, with *Lotmaria passim* BRL (2024) clustering together with the earlier assembly of *Lotmaria passim* strain SF (ATCC PRA-403) (Runckel et al. 2014) (Figure 1). *Crithidia mellificae* is one exception, with 16,228 predicted proteins (Kostygov et al. 2024), a complete single-copy BUSCO score of 64.6%, and twice as many contigs as any of the other sequenced species in the clade, suggestive of a suboptimal assembly (Figure 1).

Our assembly is larger and more complete than the earlier assembly of *Lotmaria passim* strain SF and is one of only four near-chromosome-level or chromosome-level assemblies among the 10 species and 13 individual assemblies that we compared (Figure 1). When we analyzed the completeness of our assembly using a set of 130 Euglenozoa benchmarking universal single-copy orthologs (BUSCOs) we achieved a score of 100% (Table 1), which fits well within our expectations from closely related species (90% to 100% complete single copy BUSCOs, Figure 1). Our annotation subsequently identified 128 complete single-copy BUSCOs (98.5%, Table 1, Figure 1) and 2 duplicated BUSCOs (1.5%, Table 1, Figure 2). Both duplications were found on chromosome 5 and chromosome 6 (Figure 2) and may be the result of a chromosomal duplication event, as discussed in detail below.

**Figure 2.**
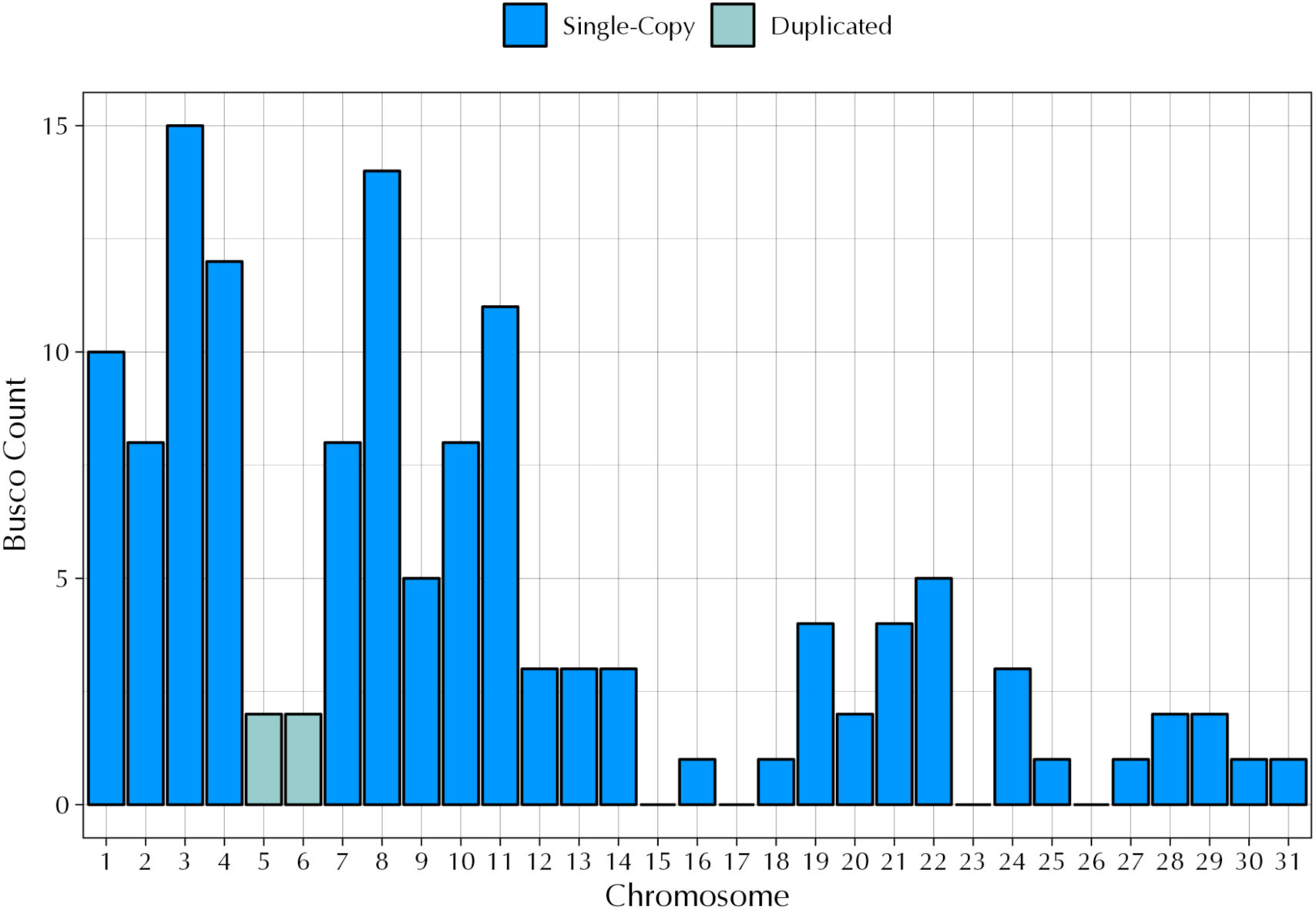
BUSCO count by chromosome. The x-axis denotes the chromosome number and the y-axis denotes the BUSCO count derived from 130 Euglenozoa benchmarking universal single-copy orthologs (BUSCOs). Of the 31 chromosomes in the *Lotmaria passim* BRL (2024) genome, chromosomes 15, 17, 23, and 26 do not contain any BUSCOs. Chromosome 5 and chromosome 6 each contain two duplicated BUSCOS, an acetyl-CoA carboxylase (OrthoDB 199at33682; v10-1.orthodb.org) and peptidase M41 (OrthoDB 5001at33682; v10-1.orthodb.org).

### Variation in Somy Within *Lotmaria passim* BRL

The expansion of genomic resources within the family Trypanosomatidae has increased exponentially in recent years, with a new focus on previously under-represented species beyond those of direct medical or veterinary importance. As more of these resources have become available, it has become abundantly clear that these genera often display varying degrees of aneuploidy (Albanaz et al. 2023), often taking the form of either—or in some cases, both—chromosome loss or duplication. The increased plasticity observed in chromosome number and adaptive changes in somy and ploidy within Trypanosomatidae is likely one of several mechanisms to control gene expression that are employed to bypass their lack of transcription-level, gene-specific regulation (Sterkers et al. 2012; Lukeš et al. 2018; Maslov et al. 2019).

The *Lotmaria passim* BRL (2024) genome is no exception and displays several cases of aneuploidy, which we inferred from a combination of sequencing depth (Figure 3, Supplementary Figure S2) and variant calling (Supplementary Figure S3). We determined that 28 of the 31 chromosomes (92% of the genome) display evidence of disomy, while only 3 chromosomes (chromosomes 9, 16, and 17; 8% of the genome) display evidence of trisomy (Figure 3). Although there are no definitively tetrasomic chromosomes present in the *Lotmaria passim* BRL (2024) assembly, there is evidence that chromosomes 5 and 6 are likely derived from a single tetrasomic chromosome which then diverged from one another in an evolutionarily recent gain in chromosome number.

**Figure 3.**
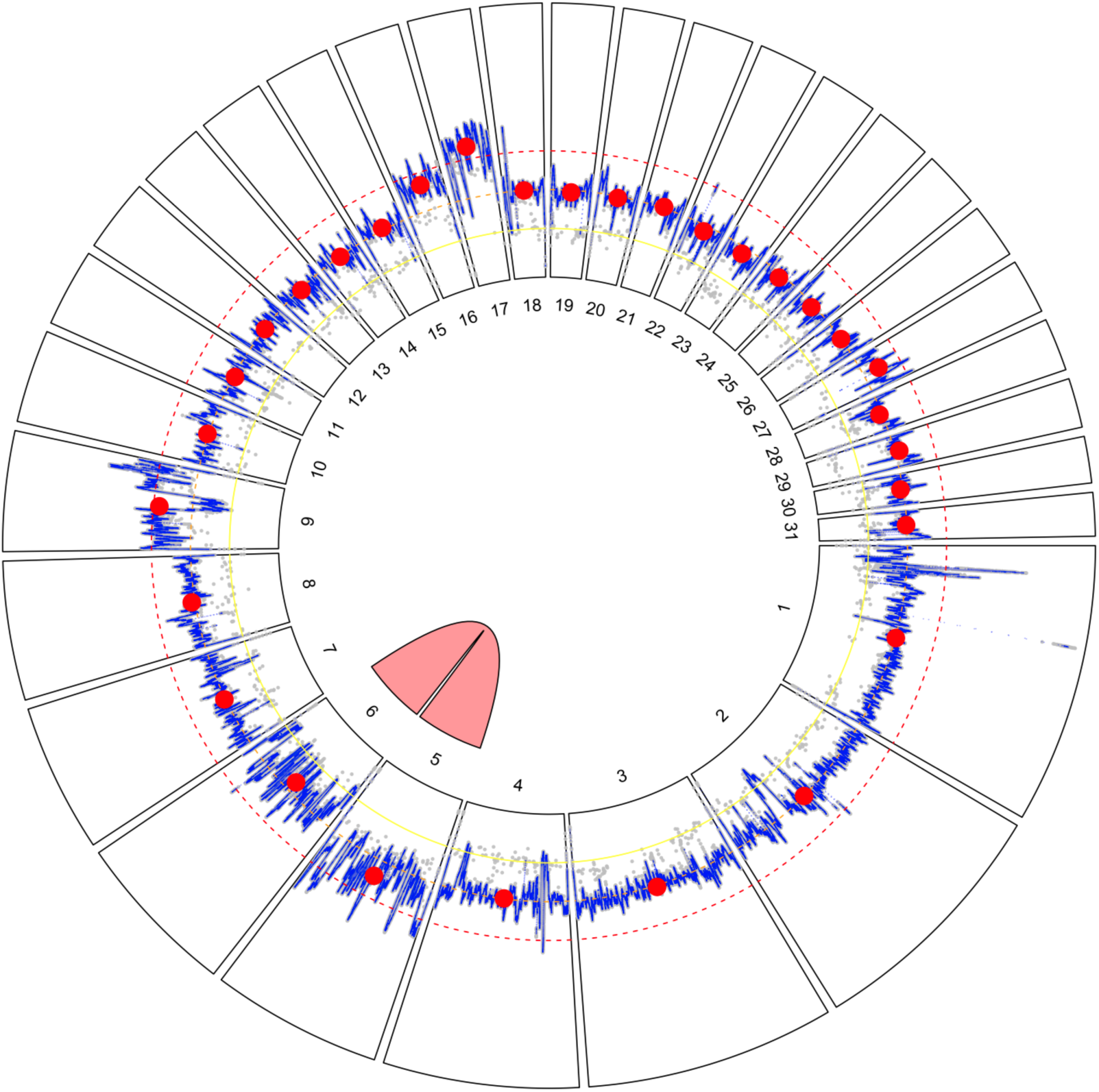
Circular plot of *Lotmaria passim* BRL (2024) genome assembly. Radial segments correspond to the 31 nuclear chromosomes and are numbered by size from largest to smallest. Sequencing depth is presented within each segment. The gray points show baselevel read depths, subsampled at 100 bp intervals, while the blue line trace represents a 1 Kb moving average, and the red circles represent chromosome-level medians. Concentric yellow, orange, and red dashed lines represent 50%, 100%, and 150% of the median coverage depth, which corresponds to the expected depths for monosomic, disomic, and trisomic chromosomes, respectively. The pink link between chromosome 5 and chromosome 6 indicates high similarity (98.62%) between these two regions, suggesting a chromosomal duplication event previously occurred. The faint scatter of gray points roughly half the read depth of the chromosomes overall (referred to as the ‘secondary trace’ in the text) suggests that heterozygous sites are conspicuously absent from chromosome 21, suggesting that our strain harbors two identical haplotypes of this chromosome.

Chromosome 5 and chromosome 6 are remarkably similar (Figure 3), and when aligned with NCBI BLAST, they have a sequence similarity score of 98.62%. Although chromosome 5 has a normalized read depth of 2.5x and chromosome 6 has a normalized read depth of 1.7x (Figure 4), both chromosomes are likely disomic but will require further study to ascertain their somy status with more certainty. The doubled normalized read depth deviations from the 2x expectation for disomic chromosomes could reflect duplicate read mapping due to the two regions’ high degree of similarity. This would complicate inferences via normalized read depth alone, particularly for chromosome 5, where the interquartile range in depth at individual bases spans most of the 2x to 3x interval (Supplementary Figure S2). When we performed variant calls on both of these chromosomes and compared our assembly against an earlier sequencing effort completed in 2016 by the University of Massachusetts (GenBank Accession ID: GCA_002216525.1), there was a very low variant count in the BRL (2024) assembly and distributions leaning towards tetrasomy in BRL (2016) (Supplementary Figure S3), likely due to difficulties in distinguishing between the two chromosomes.

**Figure 4.**
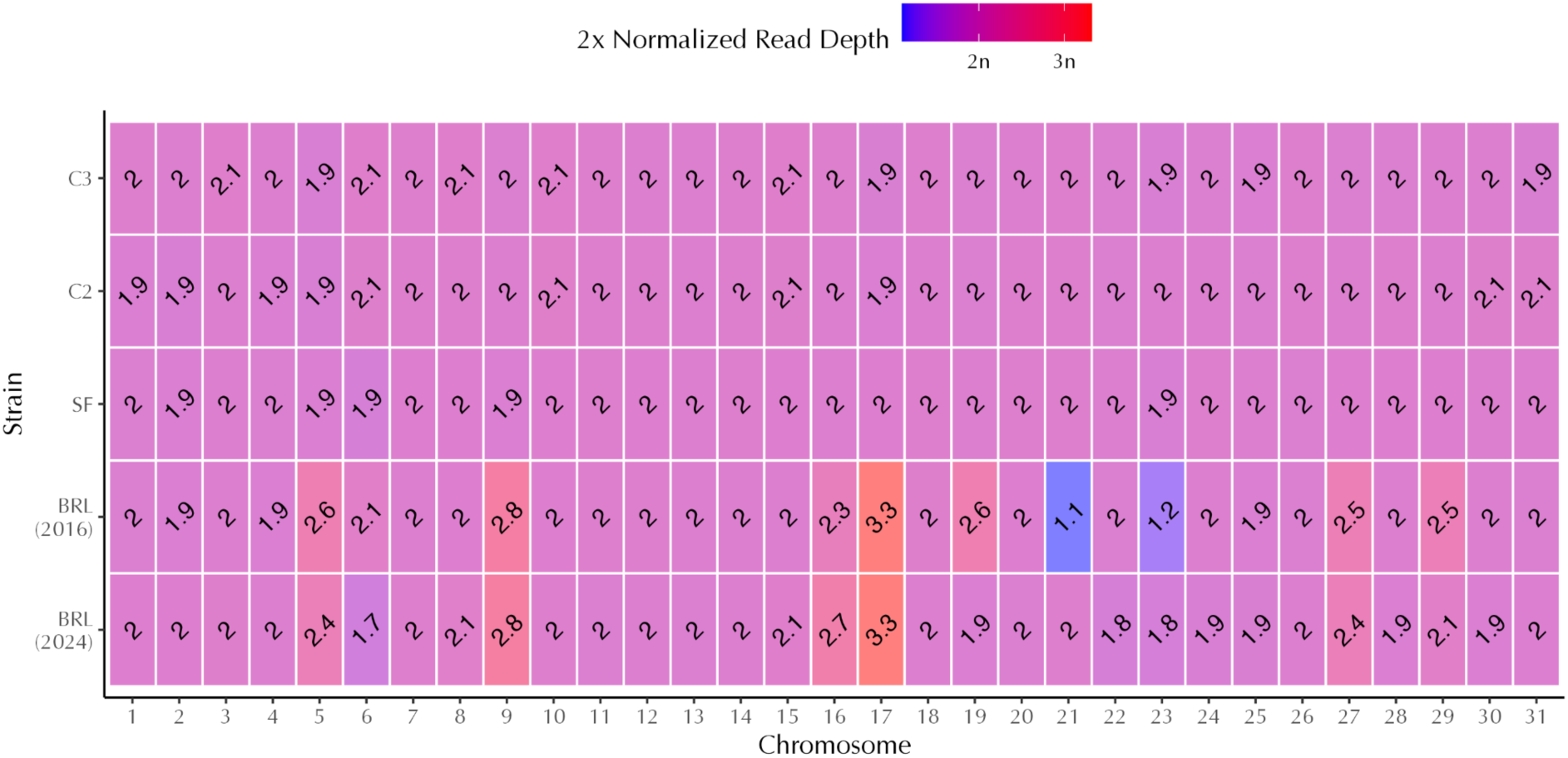
Heatmap of normalized sequencing depth for five *Lotmaria passim* genome assemblies. The x-axis denotes the nuclear chromosome number and the y-axis denotes the *Lotmaria passim* strain of origin for each assembly. Shading and text within each box depict the median sequencing depth of each chromosome when mapped to the *Lotmaria passim* BRL (2024) assembly after normalization to the median chromosome-level depth of the corresponding assembly. These depths are used as a tentative indicator of somy (i.e. chromosome copy number), with a 2x normalized read depth of 1x, 2x, and 3x denoting a monosomic, disomic, or trisomic chromosome, respectively. We display doubled normalized depth values such that the number on the plot corresponds to chromosome copy number. *Lotmaria passim* BRL (2016) normalized read depth indicates the loss of one haploid chromosome in comparison to strains C2, C3, and SF, while *Lotmaria passim* BRL (2024) normalized read depth provides evidence for the recent duplication of chromosome 21, returning it to a state of disomy in this assembly.

Sequencing depth and variant calling for strain BRL (2024) did not conform to expectations for discrete levels of somy for 6 of the 31 nuclear chromosomes. To further demystify the observed aneuploidy of *Lotmaria passim* BRL (2024), we compared our previously calculated metrics to the normalized read depth and variant calling of the BRL (2016) assembly which was completed shortly after the strain’s deposition at ATCC and lacks the approximately six months maintained in active culture (∼500 generations) and several cryopreservation events undergone in our laboratory prior to sequencing of the BRL (2024) strain (Figure 4). While this comparison revealed that the majority of the *Lotmaria passim* BRL (2024) chromosomes are indeed disomic, it also revealed several deviations from disomy.

Comparison of the two BRL assemblies identified five primary somy differences. Chromosome 16 is likely a mixture of both disomic and trisomic chromosomes, with trisomy emerging as the dominant form within strain BRL (2024), while chromosome 19 appears to be undergoing an opposite shift, with disomy evolving from trisomy (Figure 4). These shifts are further evidenced by variant calls, which lean towards trisomy and disomy, respectively (Supplementary Figure S3). Chromosome 23 is shifting towards disomy in strain BRL (2024) and is likely derived from a mixture of haploid and diploid cells, with normalized read depth elevated from 1.2x in strain BRL (2016) to 1.8x in strain BRL (2024) (Figure 4).

In another departure from clear-cut disomy, chromosome 27 is elevated above 2x normalized read depth (Figure 4) at 2.5x in BRL (2016) and 2.4x in BRL (2024). Variant calls (Supplementary Figure S3) in both strains suggest that there is a mixture of diploid and triploid cells present, likely with two identical copies and one distinct copy within the sequenced population. Although normalized read depth (Figure 4) suggested two copies per cell of chromosome 28, the secondary coverage trace (Supplementary Figure S2) exceeds 50% of the main trace, suggesting a departure from the 1:1 distribution of allelic variants typical of a diploid, and variant plots (Supplementary Figure S3) indicate close to a 2:1 ratio of alleles at heterozygous sites. This could arise if the population contains a mixture of conventional diploids, which carry two distinct copies of the chromosome, and unconventional ones that harbor two identical copies. There is strong evidence that chromosome 29 is disomic in BRL (2024) based on a doubled normalized read depth of 2.1x, although BRL (2016) leans towards trisomy at 2.5x (Figure 4). The variant calls for each of these strains provide evidence that BRL (2016) likely contained a mixture of diploid and triploid cells at the time of sequencing, while BRL (2024) has again shifted to a mixture of diploids with two distinct and two identical copies of the chromosome.

The last and arguably most remarkable evidence of *Lotmaria passim*’s plasticity in somy lies within chromosome 21. During our assembly and further analysis of the BRL (2024) reference genome, we noticed an apparent lack of the base-level read depths (secondary trace) hovering at roughly half the read depth of the chromosomes overall which are apparent in the other 30 chromosomes (gray points; Figure 3, Supplementary Figure S2). This suggests that the several heterozygous sites in each of the other 30 chromosomes in the genome are conspicuously absent in chromosome 21, leading us to believe that a single monosomic copy was recently duplicated to bring chromosome 21 to a state of disomy. We found that chromosome 21 in the BRL (2016) strain appeared to be monosomic as reflected by its 1.1x normalized read depth, while the BRL (2024) assembly appears disomic (Figure 4). We hypothesize that since the BRL (2016) assembly, the BRL isolate underwent a duplication of the remaining copy of chromosome 21 during axenic cell culture expansion, which thereby restored chromosome 21 to disomy in subsequent cell cultures that were eventually used for sequencing of the BRL (2024) strain. The uniquely low heterozygosity observed in chromosome 21 of the BRL (2024) assembly would thus be explained by such a recent duplication event and by the limited time these cultures are maintained in an active state over which unique mutations could accrue; cultures are frozen as stocks and grown only when expansions are needed. Both the 2x normalized read depth (Figure 4) and lack of variants (Supplementary Figure S3) support this hypothesis, although karyotyping will likely be necessary for unambiguous inference.

One potential explanation for these shifts within the earlier BRL (2016) and current BRL (2024) populations could be due to the intervening period during which the line has been maintained in culture, which includes approximately six months (∼500 generations) and several cryopreservation events in our laboratory, as well as shifts from antibiotic-supplemented Schneider’s medium (Schwarz et al. 2015) used during initial cultivation to modified FPFB medium (Salathé et al. 2012) which we currently use to support greater cell densities. It is widely accepted that relatively rapid changes in chromosome copy number are considered a mechanism by which trypanosomatids adapt to changing environments (Sterkers et al. 2012; Reis-Cunha et al. 2018; Albanaz et al. 2023). This adaptation is evidenced by evolution experiments in *Leishmania*, suggesting that both time and changes in growth conditions may have contributed to the inferred genomic changes (Ponte-Sucre et al. 2017). Microevolution assays performed on *Trypanosoma cruzi* also evidence these adaptations, suggesting that continuous time in culture led to ‘genome erosion’ events as tetraploid hybrids shifted towards stable populations of triploid hybrids (Matos et al. 2022).

### Variation in Somy Across Strains

To investigate patterns of aneuploidy in Lotmaria passim outside of the BRL strain, we conducted parallel analyses on three additional, independently isolated lines: a 2016 sequencing of the SF strain used for the original Lotmaria passim draft assembly (Runckel et al. 2014) and the recently sequenced Spanish isolates named ‘C2’ and ‘C3’ (Buendía-Abad et al. 2021). In contrast to the eccentricities in copy number suggested by the two assemblies of the BRL strain (2016 sequencing effort (GenBank Accession ID: GCA_002216525.1) and 2024 sequencing effort (GenBank Accession ID: GCA_037349495.1)), none of the three other strains sequenced showed convincing evidence of deviation from expectations for disomy at any chromosome. Although allele frequencies exhibited some pattern of tetrasomy for chromosomes 5 and 6, analysis sequencing depth suggested disomy (Supplementary Figure S2); the anomalous allele frequencies likely reflect difficulty distinguishing these two highly similar chromosomes using short reads. These findings leave the extent of aneuploidy in natural populations and its ramifications for parasite fitness unclear. Given that the freshly isolated, low passage C1 strain resulted in greater host mortality as opposed to the more extensively cultured strain SF in experimentally inoculated honey bees (Buendía-Abad et al. 2021), understanding how both laboratory cultivation and other environmental shifts affect the genome evolution and within-host dynamics of this parasite is warranted and has potential to elucidate mechanisms of adaptation in this host-parasite system.

Moreover, the extent of aneuploidy exhibited in the BRL strain lies well within ranges reflected by read-depth analyses of *Trypanosoma cruzi* discrete typing units TcII and TcIII (Reis-Cunha et al. 2018) and found in over twenty other trypanosomatids of varying ages post-isolation (Albanaz et al. 2023). Paired with the evidence for a past duplication of chromosomes 5 and 6, it is likely that copy numbers are flexible in naturally circulating *Lotmaria passim* populations as well. In addition, within-species differences in ploidy exist across isolates of the closely related *Leptomonas pyrrhocoris* (Flegontov et al. 2016). Sequencing of additional *Lotmaria passim* strains would be necessary to gain a more complete representation of how somy varies within and across populations, as well as its environmental correlates and consequences for parasite infectivity and reproduction.

## CONCLUSIONS

The Lotmaria passim BRL (2024) genome is one of the most comprehensive genomic resources available for monoxenous trypanosomatids within the subfamily Leishmaniinae and greatly improves upon the previously available reference assembly (Runckel et al. 2014), providing a high quality resource for future work that promises insights into the evolutionary changes leading to specialization within honey bees. Our assembly provides additional examples of aneusomies within trypanosomatids and suggests further study of the causes and consequences of variation in somy in monoxenous species is necessary. The honey bee-Lotmaria passim system offers many parallels to the challenges faced by other Leishmaniinae during infection of mammals and exposure to antiparasitic drugs and will be an important tool for understanding mechanisms of infection and adaptation in their medically important dixenous relatives.

## DATA AVAILABILITY

All *Lotmaria passim* BRL (2024) Hi-C and HiFi raw reads are available on NCBI GenBank (accession ID: SRX22798691 and SRX22798690) and the assembled chromosomes (accession ID: GCA_037349495.1), maxicircle, and minicircles are listed under BioProject PRJNA1049372. The annotated genome is available on NCBI GenBank (accession ID: [to be added]) and FigShare ([link to be added]), and detailed methods regarding both the assembly and annotation can be found herein within the manuscript. All supplementary material can be found on FigShare ([link to be added]).

## ACKNOWLEDGMENTS

The authors would like to thank Amanda Albanaz for proteome annotations of *Crithidia mellificae, Crithidia bombi,* and *Crithidia expoeki*.

## SUPPLEMENTARY INFORMATION

**Supplementary Figure S1.**
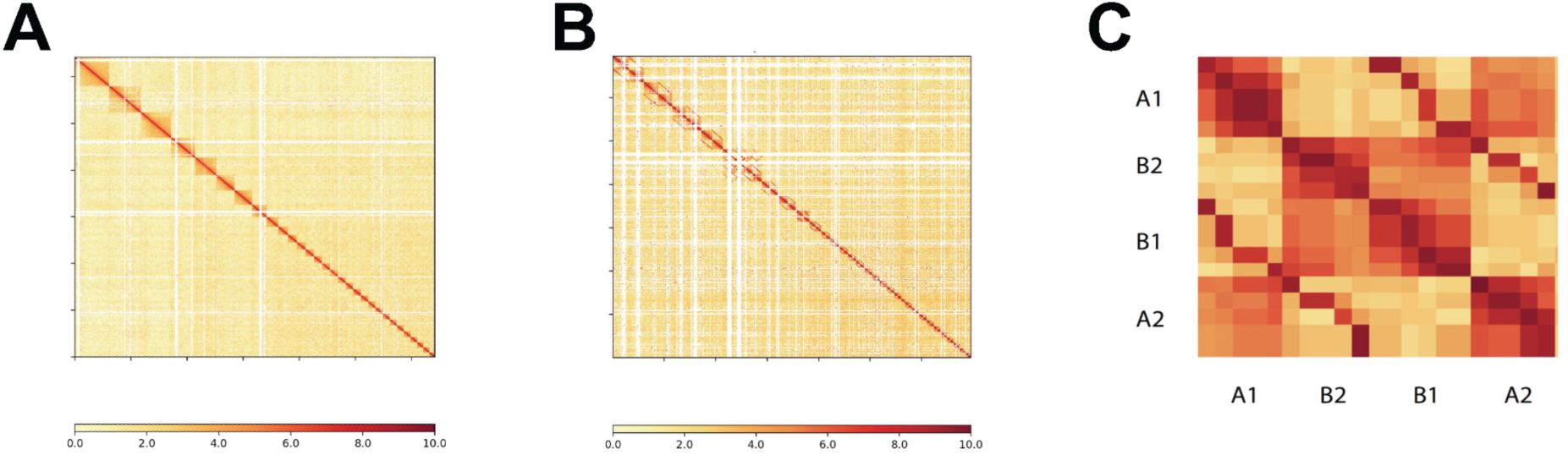
Genome Assembly Hi-C Contact Maps. Hi-C contact maps of (A) the principal nuclear genome assembly excluding chromosome 6 with 10 kb bin size, (B) the diploid nuclear genome assembly with 100 kb bin size, and (C) haplotigs of a chromosome with a switch error. Chromosomal sequences are ordered by index. In the diploid contact map (B), alternative sequences are placed next to their corresponding principal sequences. Color intensity represents the normalized number of valid Hi-C read pairs shared between bins on the x and y axes. Each chromosome forms a distinct block with higher within-chromosomal contact frequency. Off-diagonal lines in (B) and C) arise from mismapping read pairs from homologous regions between haplotigs. Despite this, blocks indicating sequence boundaries remain evident. Panel C illustrates how a switch error disrupts the top-left and bottom-right blocks and should be manually corrected. Letters represent haplotigs and numbers represent regions.

**Supplementary Figure S2.**
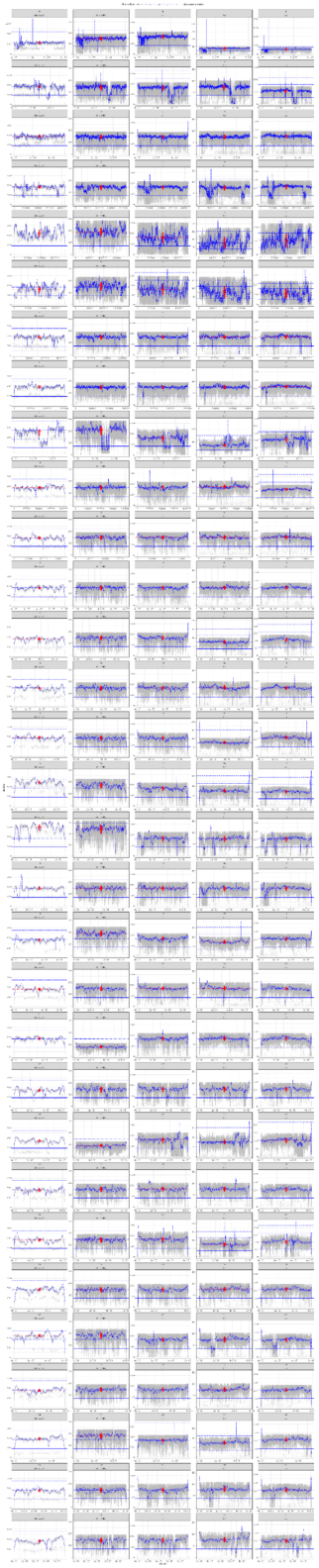
Base-level sequencing depth for each of the 31 nuclear chromosomes of *Lotmaria passim* BRL (2024) assembly. The x-axis shows the position on the chromosome and the y-axis shows the read sequencing depth. Gray dots show depth for individual positions and the blue line trace represents the median depth over a rolling 5 Kb window. The red point range at the chromosome midpoint represents the chromosome-level median and interquartile range. Horizontal blue lines represent 50%, 100%, 150%, and 200% of the median chromosome-level depth for the assembly overall which corresponds to the expected depths for monosomic, disomic, trisomic, and tetrasomic chromosomes, respectively. The red horizontal line shows the observed median. Plots include the lower 95% of data points to better visualize the 50% to 200% range of the median chromosome-level depth for the assembly overall.

**Supplementary Figure S3.**
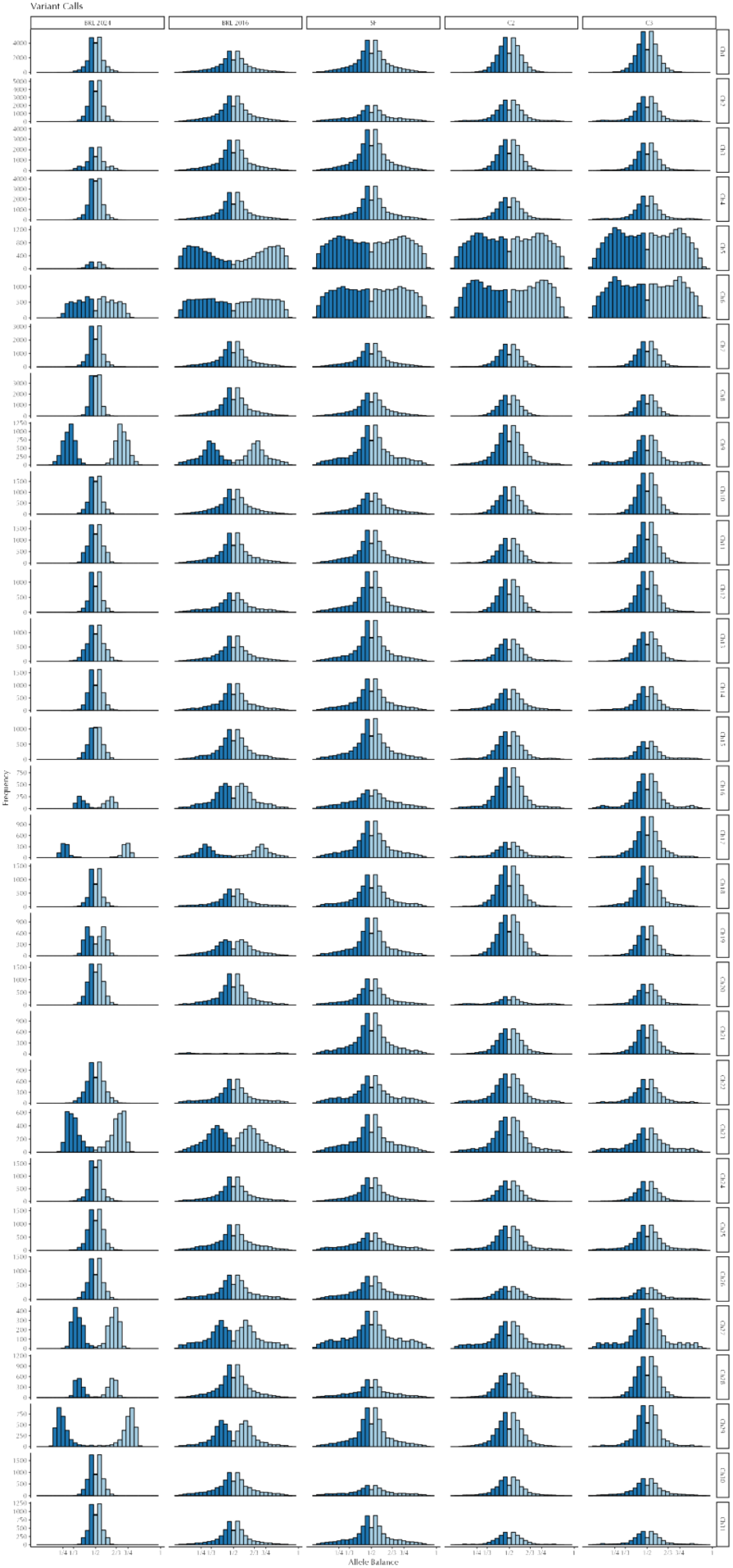
Histogram of allele frequencies at polymorphic sites for each of the 31 nuclear chromosomes estimated from reads for each of five genome sequencing projects. The expected frequency distributions for each ploidy level are two peaks centered at 1/2 for diploids; two peaks centered at 1/3 and 2/3 for triploids; and 3 peaks centered at 1/4, 1/2, and 3/4 for tetraploids. No obvious peaks are expected for haploids.

